# Effects of increasing hydrostatic pressures on marine microbial enzymatic activities

**DOI:** 10.1101/2025.01.12.632627

**Authors:** John Paul Balmonte, Carol Arnosti, Douglas H. Bartlett, Stephanie Caddell, Ronnie N. Glud

**Author notes:** **Correspondence:** John Paul Balmonte.

## Abstract

High hydrostatic pressure is characteristic of the deep ocean and presumed to influence microbial functions and viability. However, marine microbial processes are typically measured only at atmospheric pressure (0.1 MPa), limiting our understanding of pressure effects on the activities of microbes that sink as part of the biological carbon pump as well as those that reside in the deep ocean. To test pressure effects on microbial functions, we measured extracellular enzymatic activities—the first step in organic matter remineralization—of a moderate piezophile (Photobacterium profundum SS9), as well as of microbial communities in waters and sediments from shallow to abyssal (5,500 m) depths and their cell-free enzymes (<0.2 µm). Activities were measured at 0.1-100 MPa to assess the pressure effects across the nearly full range of oceanic depths. *P. profundum* SS9 enzymes show varying pressure effects, from considerable stimulation at optimal pressure (28 MPa) to near complete inhibition (100 MPa). Pressure effects were measured for diverse protein- and carbohydrate-degrading and phosphorus-acquiring enzymes among pelagic and benthic microbial communities. The most common pressure effect is partial activity reduction, indicating a dampening of the initial step of carbon remineralization at increasing pressures. However, retention of cell-free enzymatic activities at higher pressures was occasionally observed even for enzymes from surface-originating assemblages, indicating functionality down to hadal depths and their potential for piezotolerance. These variable pressure effects must be considered when quantifying degradation rates of sinking and deposited particulate matter at increasing pressures in the deep ocean.

## Introduction

Marine microbial communities can traverse large vertical depths while associated with sinking organic matter. As microorganisms are displaced vertically, they encounter increasing hydrostatic pressures, which in the deepest parts of the ocean can exceed 100 MPa (equivalent to a depth of 10,000 m). The extent to which increasing hydrostatic pressure affects microorganisms in part determines their fate and viability in the deep sea (Jannasch and Wirsen, 1984; Bartlett et al., 2007) and has consequences for the biogeochemical processes that they mediate during their descent. The ability of microorganisms to maintain their biogeochemical functions when exposed to increasing hydrostatic pressure, however, remains understudied (Jannasch and Taylor, 1984; Picard and Daniel, 2013; Tamburini et al., 2013), particularly at pressures characteristic of bathyal to hadal environments (Stief et al., 2021; Amano et al., 2022; Stief et al., 2023).

Several pathways deliver microbial cells and organic matter from the surface to the deep ocean. As part of the biological carbon pump (Iversen, 2022) and particle injection pump mechanisms (Boyd et al., 2019), ca. 6-10 Pg of C is exported out of the upper water column (Burd et al., 2010), delivering up to 10^10^ bacterial cells associated with sedimenting particles to the deep sea (Turley and Mackie, 1995). These particles may be part of the elevated carbon flux following phytoplankton blooms (Poff et al., 2021), which add to the flux of organic carbon reaching the sediments. Sediment carbon accumulation may also be due to mass wasting events, which can increase sediment deposition by about a factor of 4, over timescales of ca. 150 years (Oguri et al., 2022). The deposition of organic carbon in sediments implies that a portion of surface-derived organic carbon remains unused during its downward transport (Danovaro et al., 2003; Zabel et al., 2022). This organic carbon fuels high rates of microbial oxygen demand, as observed in hadal sediments, hinting at its labile nature (Glud et al., 2013; 2021). However, the factors that prevent full remineralization of bioavailable organic carbon during descent require further study.

One hypothesis is that hydrostatic pressure, particularly at moderate (40-60 MPa) to high levels (>60 MPa), may reduce or fully inhibit microbial activities necessary to remineralize organic matter during its downward transport (Jannasch and Wirsen, 1984). Pressure-related reduction of microbial activities can occur at multiple steps of organic matter remineralization (Jannasch and Taylor, 1984; Picard and Daniel, 2013; Tamburini et al., 2013). Aerobic respiration of organic carbon by surface-originating microorganisms is partially inhibited from 10-50 MPa, but fully inhibited from 60-100 MPa (Stief et al., 2021; 2023). Even microbes in the deep sea may be affected by high hydrostatic pressure: uptake of leucine to produce microbial biomass at 40 MPa (4000 m) in the bathypelagic operates at ca. 33% efficiency when compared to uptake at atmospheric pressures (Amano et al., 2022).

High hydrostatic pressure may also decrease the efficiency of the initial degradation step of high molecular weight organic matter, which is catalyzed by diverse extracellular enzymes (Arnosti, 2011). These enzymes differ in terms of the macromolecules hydrolyzed (e.g., proteins and polysaccharides), modes of action (terminal unit-cleaving exo-acting enzymes, vs. midchain-cleaving endo-acting enzymes), and the specific structures that they target. To date, most investigations of pressure effects on extracellular enzymes have focused primarily on the activities of leucine aminopeptidase (e.g. Tamburini et al. 2002; 2003; 2009), including an investigation of upper ocean microbial communities that showed that leucine aminopeptidase activities are reduced up to 5-fold when exposed to gradually increasing pressures (Tamburini et al., 2006). In addition, a recent investigation of a range of polysaccharide hydrolase activities in coastal seawater showed partial reduction of activity at 20 MPa, and complete inhibition of activity at a pressure of 40 MPa (Lloyd et al., *in revision*). Through partial reduction or complete inhibition of extracellular enzymatic activities, hydrostatic pressure thus may help preserve sinking organic matter during its vertical descent. However, there are considerable gaps in our understanding of pressure effects on microorganisms from distinct habitats, the differential effects on activities of diverse hydrolytic enzymes, and the pressure levels at which they become inhibited. Investigating pressure effects on a wide range of hydrolytic enzymes is necessary since the enzymatic repertoire of microbial communities are diverse and distinct across regions and depths in the ocean (Baltar et al., 2010; Arnosti et al., 2011; Balmonte et al., 2021; Lloyd et al., 2022).

Whether the pressure sensitivity of microbes in the water column differs from pressure sensitivity of sediment microbial communities is also unclear. Prior studies suggest pressure sensitivity differences among microorganisms in more nutrient-rich versus oligotrophic deep-sea environments (Jannasch and Taylor, 1984), which may extend to the differences observed in sediments compared to the water column. Microbial community composition (Zinger et al. 2011) also differs significantly between the water column and sediments, as do the enzyme activities of these communities (Arnosti 2008; Teske et al. 2011). To date, however, very few studies have examined pressure effects on enzyme activities in sediments. One of these studies—which measured leucine aminopeptidase, plus α- and β-glucosidase, esterase, and chitinase—showed that enzymatic activities in deep sea sediments of the eastern North Atlantic show considerable resistance to increasing pressure levels (Poremba, 1995). An investigation of pressure effects on polysaccharide hydrolase activities in shallow coastal sediments likewise found considerable resistance to hydrostatic pressure at 20 MPa and 40 MPa (Lloyd et al., *in revision*). Preliminary evidence thus suggests that there may be fundamental differences in pressure responses and strategies between seawater and sedimentary microbial communities. Finally, the mechanisms by which hydrostatic pressure affects enzymatic activities, through potential changes in enzyme conformations that alter maximal rates and stability (Ohmae et al., 2007; Ohmae et al., 2013), changes in enzyme production (Vezzi et al., 2005), or differences in the relative distribution of specific enzymes – or the extent of functional redundancy – among community members are insufficiently known in marine systems.

Here, we investigate the effect of hydrostatic pressure on extracellular enzymatic activities, a critical function for microbial communities associated with sinking particles, microorganisms in the deep sea, as well as those in sediments that are displaced and deposited to deeper settings through offshore transport or mass wasting events. Across all sites and samples, we measured leucine aminopeptidase, beta-glucosidase, and chitinase activities; activities of endo-acting peptidases (e.g., chymotrypsins and trypsins) and phosphatase were only measured at sites in the North Atlantic. We define a pressure effect as the percentage of activity at pressures >0.1 MPa relative to that at 0.1 MPa (atmospheric pressure). We measured rates of protein- and carbohydrate-hydrolyzing and phosphorus-acquiring enzymes of a moderate piezophile—*Photobacterium profundum* SS9 (Nogi et al., 1998)—at pressures of 0.1 MPa (atmospheric pressure), 28 MPa (the optimal pressure for *P. profundum* SS9), 50 MPa, 75 MPa, and 100 MPa to identify the range of pressure effects on a bacterial isolate. Then, we applied the same experimental approach to measure pressure effects on the enzymatic activities of natural microbial assemblages from the water column and sediments. Regardless of sample depth of origin, the pressure levels used for these experiments were 0.1 MPa, 25 MPa, 50 MPa, 75 MPa, and occasionally 100 MPa, reaching pressure levels encountered in hadal environments. A consistent matrix enables the comparison of performance by different microbial communities at increasing pressures. In addition, changes in enzyme activities due to exposure to high hydrostatic pressure reflects the responses of mixed and diverse microbial cells, as well as changes in the catalytic efficiency of their wide range of cell-free enzymes. To distinguish between these possibilities, we then compared enzyme activities of unfiltered (cell-present) and 0.2 um-filtered (cell-free) seawater following exposure to particulate organic matter (POM) sources, which is intended to stimulate the production of freely-released enzymes. Determining these potential mechanisms is ecologically and biogeochemically important, as the enzymatic degradation of organic matter can happen outside the cell by enzymes released from producer cells (Vetter & Deming 1999; Traving et al., 2015) or by enzymes that remain attached to the cell surface (Reintjes et al., 2017). Testing the effects of high hydrostatic pressure on marine microbes expands our understanding of the ecology of sinking and deep-sea microorganisms, as well as the biogeochemical processes that sinking microorganisms can continue to mediate at increasing depths.

## Materials and Methods

### Site description and sample collection

Seawater and/or sediment samples were collected from several sites in Aarhus Bay, Fram Strait, Skagerrak Strait, and the North Atlantic Ocean between 2021 and 2022 (Table 1). Aarhus Bay is located east of Aarhus, Denmark. Skagerrak Strait lies in between Denmark, Sweden, and Norway and connects North Sea to the Kattegat Sea. Fram Strait connects the Arctic Ocean (north) to the Greenland and Norwegian Seas (south) and is bound by Greenland (west) and Svalbard (east); samples in the Fram Strait were collected during the PS126 cruise on the R/V *Polarstern*. Seawater and sediment samples from the North Atlantic come from stations 21, 22, and 23 during the EN683 cruise aboard the R/V *Endeavor*.

**Table 1.**
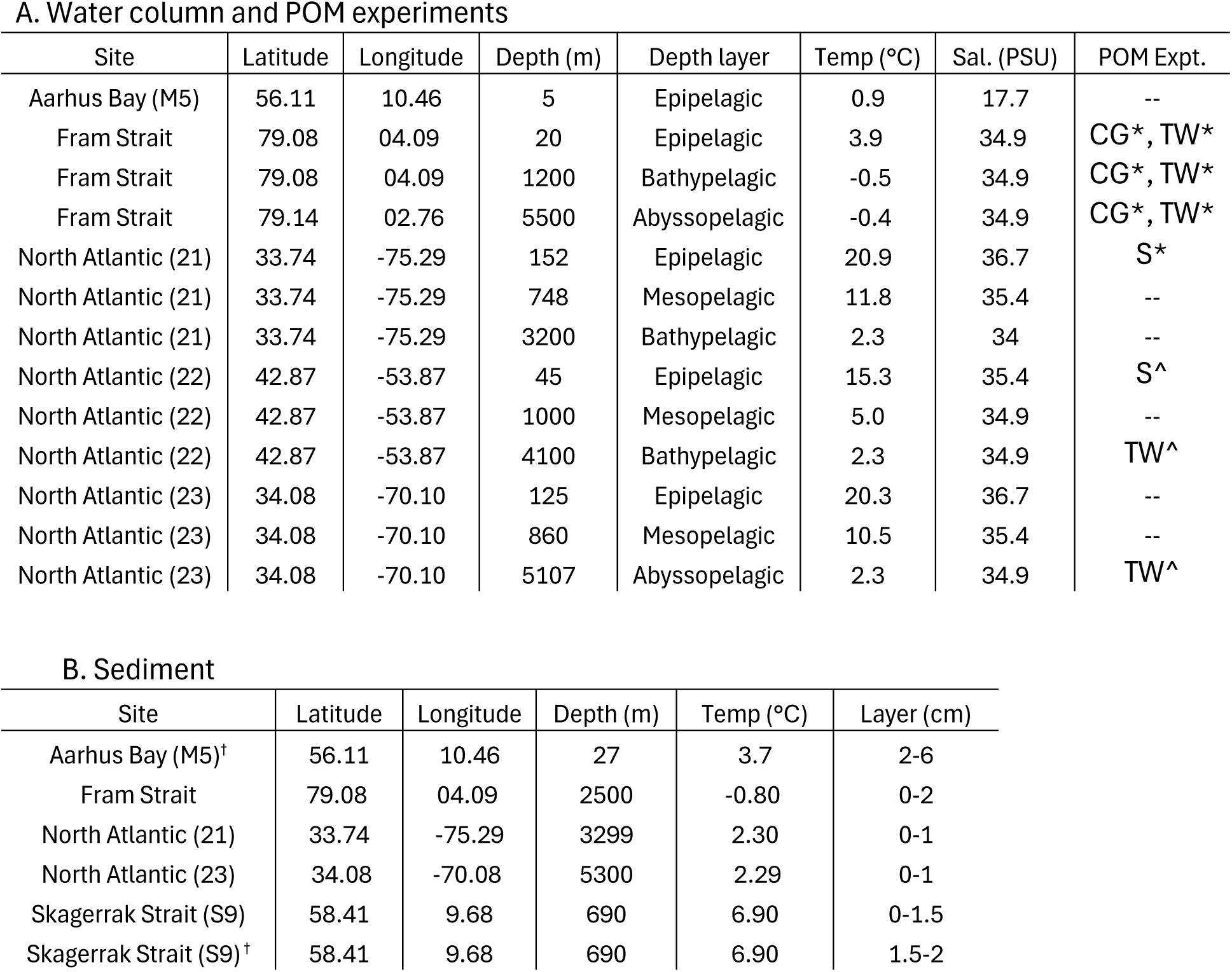
Sampling locations and physicochemical parameters for the water (A) and sediment (B) samples used to quantify pressure effects on enzyme activities. Epipelagic (≤ 200 m) were grouped separately from mesopelagic (≤ 1000 m), but bathypelagic and abyssopelagic were combined (≤ 6,000 m). Enzyme stimulation experiments with particulate organic matter (POM) marked with an asterisk (*) are those with only cell-free enzyme activity measurements, whereas those with a caret symbol (^) included a comparison of cell-free and cell-present enzyme activities. (--) denotes that no POM stimulation experiments were carried out. Sediment samples (B) marked with a cross (^†^) were collected below the oxygen penetration depth, and these experiments were carried out under anoxic conditions. All other sediment experiments used surficial sediments and were carried out under oxic conditions.

A. Water column and POM experiments
| Site | Latitude | Longitude | Depth (m) | Depth layer | Temp (°C) | Sal. (PSU) | POM Expt. |
| --- | --- | --- | --- | --- | --- | --- | --- |
| Aarhus Bay (M5) | 56.11 | 10.46 | 5 | Epipelagic | 0.9 | 17.7 | -- |
| Fram Strait | 79.08 | 04.09 | 20 | Epipelagic | 3.9 | 34.9 | CG*, TW* |
| Fram Strait | 79.08 | 04.09 | 1200 | Bathypelagic | -0.5 | 34.9 | CG*, TW* |
| Fram Strait | 79.14 | 02.76 | 5500 | Abyssopelagic | -0.4 | 34.9 | CG*, TW* |
| North Atlantic (21) | 33.74 | -75.29 | 152 | Epipelagic | 20.9 | 36.7 | S* |
| North Atlantic (21) | 33.74 | -75.29 | 748 | Mesopelagic | 11.8 | 35.4 | -- |
| North Atlantic (21) | 33.74 | -75.29 | 3200 | Bathypelagic | 2.3 | 34 | -- |
| North Atlantic (22) | 42.87 | -53.87 | 45 | Epipelagic | 15.3 | 35.4 | S^ |
| North Atlantic (22) | 42.87 | -53.87 | 1000 | Mesopelagic | 5.0 | 34.9 | -- |
| North Atlantic (22) | 42.87 | -53.87 | 4100 | Bathypelagic | 2.3 | 34.9 | TW^ |
| North Atlantic (23) | 34.08 | -70.10 | 125 | Epipelagic | 20.3 | 36.7 | -- |
| North Atlantic (23) | 34.08 | -70.10 | 860 | Mesopelagic | 10.5 | 35.4 | -- |
| North Atlantic (23) | 34.08 | -70.10 | 5107 | Abyssopelagic | 2.3 | 34.9 | TW^ |

**Table 1.** Sampling locations and physicochemical parameters for the water (A) and sediment (B) samples used to quantify pressure effects on enzyme activities.
| Site | Latitude | Longitude | Depth (m) | Temp (°C) | Layer (cm) |
| --- | --- | --- | --- | --- | --- |
| Aarhus Bay (M5)<sup>†</sup> | 56.11 | 10.46 | 27 | 3.7 | 2-6 |
| Fram Strait | 79.08 | 04.09 | 2500 | -0.80 | 0-2 |
| North Atlantic (21) | 33.74 | -75.29 | 3299 | 2.30 | 0-1 |
| North Atlantic (23) | 34.08 | -70.08 | 5300 | 2.29 | 0-1 |
| Skagerrak Strait (S9) | 58.41 | 9.68 | 690 | 6.90 | 0-1.5 |
| Skagerrak Strait (S9)<sup>†</sup> | 58.41 | 9.68 | 690 | 6.90 | 1.5-2 |

### Substrates for enzyme activity assays

The activities of various peptide- and glucose- hydrolyzing, and phosphate-acquiring enzymes were measured across a range of hydrostatic pressures up to 100 MPa, equivalent to hydrostatic pressures at an oceanic depth of 10,000 m. Enzyme assays were carried out using substrates labelled with fluorophores, either methylumbelliferyl (MUF) or methylcoumarin (MCA) (Hoppe, 1983). MUF-labelled compounds include β-glucopyranoside (β-glu; to measure beta-glucosidase), phosphate (to measure phosphatase), and N-acetyl-glucosaminide (NAG; to measure chitinase). MCA-labelled substrates include leucine (Leu; to measure leucine aminopeptidase), alanine-alanine-phenylalanine (AAF; to measure chymotrypsin), alanine-alanine-proline-phenylalanine (AAPF; to measure chymotrypsin), Boc-glutamine-alanine-arginine (QAR; to measure trypsin), and phenylalanine-serine-arginine (FSR; to measure trypsin). Enzyme activities using these substrates have been detected from surface to deep waters, particles, and sediments (Baltar et al., 2009; Balmonte et al., 2018; Balmonte et al., 2021; Lloyd et al., 2022), but only a limited few have been measured at increased hydrostatic pressures (Tamburini et al., 2002; Tamburini et al., 2006; Tamburini et al., 2009), especially at levels typical of the bathypelagic realm and below (Poremba, 1995).

### Pure culture enzyme assays

*P. profundum* SS9 was grown from −80°C cryopreserved samples in glycerol. *P. profundum* SS9 cells were streaked on a 15% agar plate containing marine broth (MB) 2216 medium (Becton Dickson Difco) and grown for 1 week at 15°C. Colonies were subsequently picked and grown in heat-sealed Pasteur-pipette plastic bulbs containing sterile MB 2216 with 0.05 M glucose and 0.1 M HEPES buffer; bulbs were incubated at 28 MPa in pressure vessels and 15°C for 3 d. After 3 d of incubation in pressure vessels, *P. profundum* SS9 cultures were dispensed from the bulbs (ca. 5 mL) and added to 1 L of sterile MB 2216. This sample (*P. profundum* SS9 + MB 2216) was used for enzyme assays immediately after mixture as described for seawater enzyme assays, but substrates were added to a final concentration of 200 µM, and the temperature used was 15°C, the optimal growth temperature for the pure culture. Tested pressures include 0.1 (atmospheric control) MPa, 28 MPa, 50 MPa, 75 MPa, and 100 MPa.

### Seawater enzyme assays

To measure enzyme activities in seawater using MCA or MUF substrates, samples were incubated in 3 mL exetainer vials amended with one substrate to a final concentration of 100 µM, which has been used as a substrate-saturating concentration across a wide range of waters (Balmonte et al., 2021). One set of enzyme assays typically consisted of triplicate live samples and one killed control using autoclaved sample water for each substrate, one live blank (no substrate addition), and one killed control blank. Enzyme activities using Leu, β-glu, and NAG were measured across all site and samples, but the remaining substrates were only used at a few sites in the North Atlantic. Regardless of sample depth of origin, a set of enzyme assays were prepared for pressure levels of 0.1 (atmospheric control), 25, 50, 75 MPa, and occasionally 100 MPa. Assays were incubated at ca. 4°C in the dark for 24 h in various types of small (Yayanos et al., 1995) and large volume pressure-retaining vessels (Stief et al., 2021; Stief et al., 2023). At the end of the 24 hr incubations, vessels were depressurized, and seawater (or cultures) in exetainers were sampled. Fluorescence was measured from 1 mL of sample and 1 mL of artificial seawater in a cuvette using a QuantiFluor mini fluorimeter. Fluorescence was converted to substrate concentration using a standard curve made of known concentrations of pure MCA or MUF fluorophore. Fluorescence measurements at t0 (upon substrate addition) were made on a replicate set of enzyme assays because measured t0 subsamples cannot be taken from the incubated exetainer vials since they need to be full for pressurization.

### Sediment enzyme assays

Enzyme activities in sediments were measured in slurries amended with either MCA or MUF substrates added to a final concentration of 200 µM. Oxic sediment slurries were made using autoclaved bottom water mixed with the oxic sediment layer at a 1:6 vol/vol dilution. In the North Atlantic sediment experiments—for which the oxygen penetration depth was not available—the upper 2 cm was used. To ensure fully oxic conditions, slurries were stirred or manually shaken and aerated for at least 2 h prior to substrate addition and start of incubation. For anoxic enzyme assays using sediments from the Skagerrak Strait, sediment slurries were made using N_2_-sparged autoclaved bottom water; slurries were further sparged with N_2_ for an additional 20 mins, anoxically dispensed in 3 mL exetainer vials, and left in the dark at 4°C to ensure anoxic conditions. Enzyme assays with oxic and anoxic sediment slurries included sorption controls set to account for substrate sorption to sediment particles and/or fluorescence quenching. Sorption controls consisted of sediment slurries amended either with MCA or MUF fluorophores at one to three known concentrations. At the end of incubations, vessels were de-pressurized to measure sample fluorescence. A subsample of 1 mL sediment slurry was added to 1 mL of artificial seawater in a microcentrifuge tube. Samples were centrifuged at 10,000 rpm for 1 min, and 1 mL of supernatant was subsampled and added to a cuvette containing 1 mL of artificial seawater. Fluorescence was measured using a mini fluorimeter, and samples were further diluted and re-measured if fluorescence values exceeded the linear range based on standard curves. Similar to seawater experiments, fluorescence measurements at t0 (upon substrate addition) were also made on a replicate set of enzyme assays.

### Cell-free vs. cell-present enzyme assays

To test the effect of pressure in the absence of cellular responses and compensation, experiments were carried out using cell-free (i.e., enzymes in the dissolved fraction after passing through a 0.2 µm pore-size filter). Since the concentrations of cell-free enzymes are often relatively low in seawater, we stimulated microbial production of enzymes by adding several types of particulate organic matter (POM): either high molecular weight *Thalassiosira weissflogii* biomass (Balmonte et al., 2019; Brown et al., 2022) and pieces of freshly collected sargassum to represent marine sources of POM (Supporting Information, Fig. S3, S5), as well as a mixture of dried cereal grasses to present terrestrial sources of POM (Supporting Information, Fig. S4). POM was added at different concentrations: 25 mg L^-1^ for the *T. weissflogii* biomass based on prior studies (Balmonte et al., 2019; Brown et al., 2022); 20 g L^-1^ for cereal grass based on tests in the lab; pieces of sargassum of unknown weight due to logistical constraints on the cruise (but see Supporting Information, Fig. S5 for the sargassum incubations). Enzyme stimulation with sargassum proceeded for 1-3 days, and 6 days for *T. weissflogii* biomass and cereal grass at atmospheric pressure and 4 °C in the dark. The different POM concentrations and incubation lengths varied to ensure or maximize the potential for enzyme stimulation. After the enzyme stimulation incubations, cells and other constituents >0.2 µm in diameter were removed using vacuum filtration through a 0.2 µm pore size 47 mm diameter cellulose membrane filter (Whatman) or using a 0.2 µm sterile cellulose acetate syringe filter. Enzyme assay setup with the cell-free enzymes followed procedures for the seawater enzyme assays. In experiments using North Atlantic seawater, cell-free enzyme activities were compared to cell-present enzyme activities in seawater enriched with *T. weissflogii* biomass and sargassum, but which were only pre-filtered using a 100 µm mesh sieve. Enzyme activity assays were identical for cell-free and cell-present experiments.

### Calculation of pressure effects

The pressure effect (PE) represents the enzyme activity rate at hydrostatic pressures above 0.1 MPa as a fraction of that measured at atmospheric pressure. The following equation was used: PE = (Rate_>0.1 MPa_ / Rate_0.1 MPa_) x 100%

## Results and Discussion

### Pressure effects on a moderate piezophile

Based on experiments that expose a moderate piezophile (*Photobacterium profundum* SS9) to pressures of 0.1-100 MPa, we observe a wide range of pressure effects on a single microorganism, from considerable stimulation of enzyme activities to almost full inhibition (Fig. 1). At its optimal pressure of 28 MPa, *P. profundum* SS9 shows a nearly 400% stimulation of leucine aminopeptidase activities relative to that at atmospheric pressure (0.1 MPa). Other enzyme activities, including chitinase (NAG; abbreviation of the chitinase substrate N-acetylglucosamine) and phosphatase are also stimulated at 28 MPa but to a significantly lesser extent—ca. 200% and 160% relative to 0.1 MPa, respectively. A near doubling of hydrostatic pressure, up to 50 MPa, reduced leucine aminopeptidase activities to only 300% stimulation relative to atmospheric pressure. Reduction of phosphatase activities from 28 to 50 MPa is proportionally less than that for leucine aminopeptidase. In contrast, NAG chitinase activities remain relatively unchanged from 28 to 50 MPa. Increasing pressure levels up to 75 MPa and 100 MPa resulted in near total inhibition of leucine aminopeptidase and phosphatase activities but shows the persistence of chitinase activities at ca. 50% relative to rates at 0.1 MPa.

**Figure 1.**
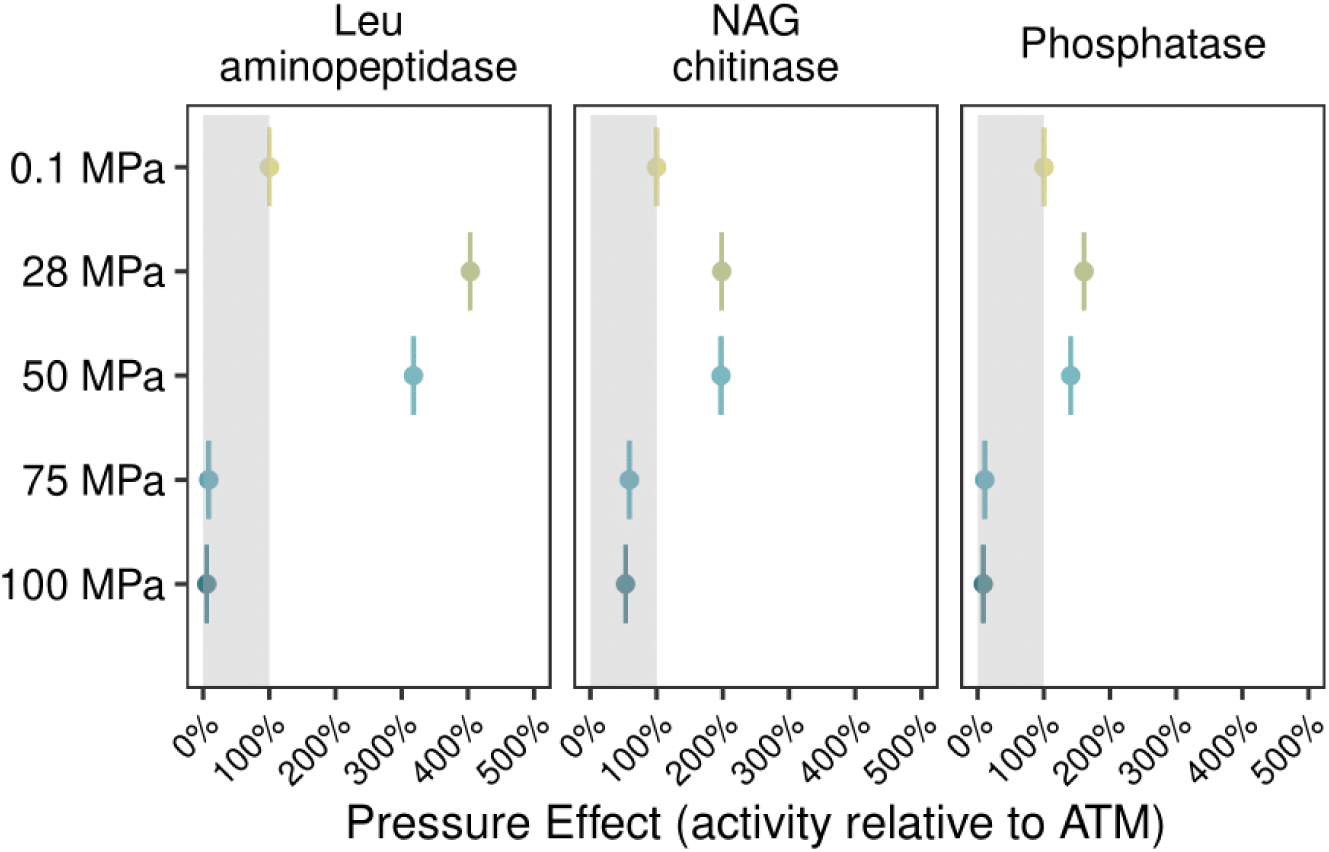
Pressure effect on enzyme activities of the moderate piezophile *Photobacterium profundum SS9*. The shaded box indicates the region of pressure reduction. Leu = leucine; NAG = N-acetyl-glucosaminide (substrate for chitinase). ATM = atmospheric pressure (0.1 MPa)

The pressure responses of *P. profundum* SS9 enzymatic activities demonstrate several important points. First, the highest enzyme activities are observed at the organism’s optimal growth pressure of 28 MPa (Bartlett et al., 2007). As cells multiply rapidly at 28 MPa, energy is invested in the production of hydrolytic enzymes to meet metabolic demands. However, on a per-cell basis, enzyme activities decrease with increasing pressure (Supporting Information, Fig. S1), indicating that production of specific hydrolytic enzymes may be increasingly challenging, or that the enzymes produced have lower catalytic efficiency at higher pressures. As observed in a different study, increased transcription of genes to produce pullulanase—an enzyme that hydrolyzes the polysaccharide pullulan—in *P. profundum* SS9 grown at 28 MPa (Vezzi et al., 2005) illustrates that pressure-regulated transcription of genes for hydrolytic enzymes is fine-tuned (Campanaro et al., 2012): while some genes for hydrolytic enzymes may be upregulated in response to increased pressure, others may be downregulated even at an organism’s optimal pressure. In addition, at pressures >50 MPa, *P. profundum* SS9 enzymatic activities are drastically reduced (Fig. 1), although the relative decrease varies across enzymes. As a caveat, the enzyme activities by *P. profundum* SS9 are profoundly affected by the culturing conditions; the addition of glucose, for example, may have had consequences on the measured pressure sensitivity of the enzyme activities. Nevertheless, these results demonstrate that enzymatic activities—and likely microbial enzymatic production—can proceed well above and below an organism’s optimal pressure. These findings additionally warrant an investigation of the pressure responses of vertically-displaced microorganisms, in sediments via resuspension and mass wasting events and in the water column through the biological carbon pump.

### Pressure effects on complex microbial assemblages

Among complex benthic microorganisms from coastal to abyssal sediments (Table 1), limited pressure reduction of enzymatic activities is observed at 25 MPa, but the most common pressure response from 50-75 MPa is partial activity reduction (Fig. 2). These results are consistent with the findings of pressure-tolerant (non-piezophilic) microorganisms in the ocean that can withstand pressures of 20-30 MPa in the ocean (Jannasch and Wirsen, 1984). The average activities relative to that at 0.1 MPa for leucine aminopeptidase, beta-glucosidase, and NAG chitinase at 25 MPa is 93%,100%, and 100%, respectively. The tight range of pressure responses across the three enzyme activities at 25 MPa is remarkable since these benthic communities come from—and are presumably adapted to—very different conditions (i.e., surface oxic and subsurface anoxic layers) and hydrostatic pressures, from coastal to bathyal sediments. In addition, evidence for pressure insensitivity or even minor pressure stimulation up to 75 MPa comes from beta-glucosidase activities collected at ca. 17 m at Aarhus Bay (Fig. 2; S2). Consistent with these findings, a recent investigation of polysaccharide hydrolase activities in coastal sediments from Denmark likewise shows considerable resistance to effects of increased hydrostatic pressure, with sedimentary activities showing only small reductions at 20 MPa (Lloyd et al., *in revision*). Our findings indicate that over the short term (24 h of pressurization), some hydrolytic enzymes from coastal benthic microbial communities can tolerate pressures of up to two orders of magnitude greater than the pressure at their depth of origin. Observations that leucine aminopeptidase activities at 75 MPa are ca. 90% relative to 0.1 MPa (Fig. 2) also demonstrate less pronounced pressure reduction of some enzymatic activities by benthic microbial communities. Previous findings of limited pressure reduction of a few enzyme activities in deep ocean sediments incubated at in situ pressures of ca. 40 MPa (Poremba, 1995) are also consistent with our observations. The high species richness of benthic microbial communities, coupled with functional redundancy of enzyme activities among these microorganisms (i.e., overlap in function across distinct taxa), may explain why specific enzymatic activities are retained despite increasing hydrostatic pressures. Consequently, the greater enzymatic capabilities of benthic microbial communities—or a high production of cell-free enzymes by benthic microbes (Schmidt et al., 2021)—may increase the possibility that some of these enzymes retain activity at high pressure, even if a subset of enzymes are inhibited by pressure.

**Figure 2.**
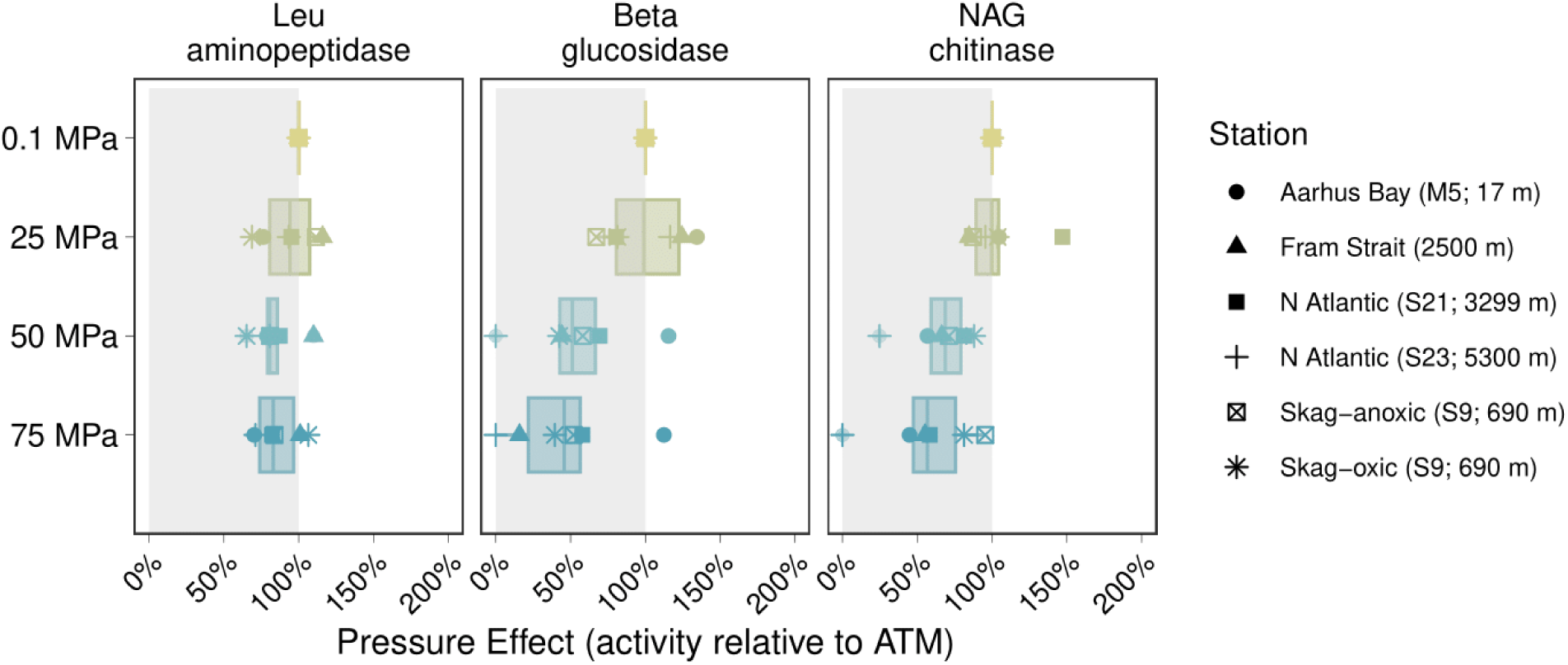
Pressure effects on enzymatic activities of natural benthic microbial communities from coastal to abyssal sediments. The shaded box indicates the region of pressure reduction. Leu = leucine; NAG = N-acetyl-glucosaminide (substrate for chitinase). Skag-anoxic = Skagerrak Strait (S9) anoxic; Skag-oxic = Skagerrak Strait (S9) oxic (see Table 1). ATM = atmospheric pressure (0.1 MPa)

To quantify pressure effects of endo-acting peptidase enzymes (e.g., AAF- and AAPF chymotrypsin, QAR- and FSR-trypsin) and chitinase in the water column, we also measured activities and pressure effects using five additional substrates in recovered water samples (Fig. 3). These enzyme activities were generally low—especially in mesopelagic and bathypelagic depths—even at atmospheric pressure; thus, we report these directly as rates and did not calculate a pressure effect. Nevertheless, enzyme activities by pelagic microbial communities showed widely varying pressure responses. With higher rates at 0.1 MPa, pressure reduction was evident at pressures >0.1 MPa, as seen for leucine aminopeptidase in Aarhus Bay epipelagic waters (Fig. 3). We interpret these varied pressure effects as a reflection of the aggregate responses of diverse enzymes that can be produced by distinct microbial communities across a range and depth of environments (Zinger et al. 2011;; Lloyd et al., 2022). Even closely-related microorganisms can differ markedly in their enzymatic capabilities (Zimmerman et al., 2013; Avcı et al. 2020). In addition, hydrolysis rates may be the result of a few highly efficient enzymes that are the product of a few select organisms, or may reflect the activities of a large range of distinctly different enzymes with comparatively low activities produced by many different community members. All of these enzymes (and their producer organisms) likewise may exhibit different sensitivities to pressure.

**Figure 3.**
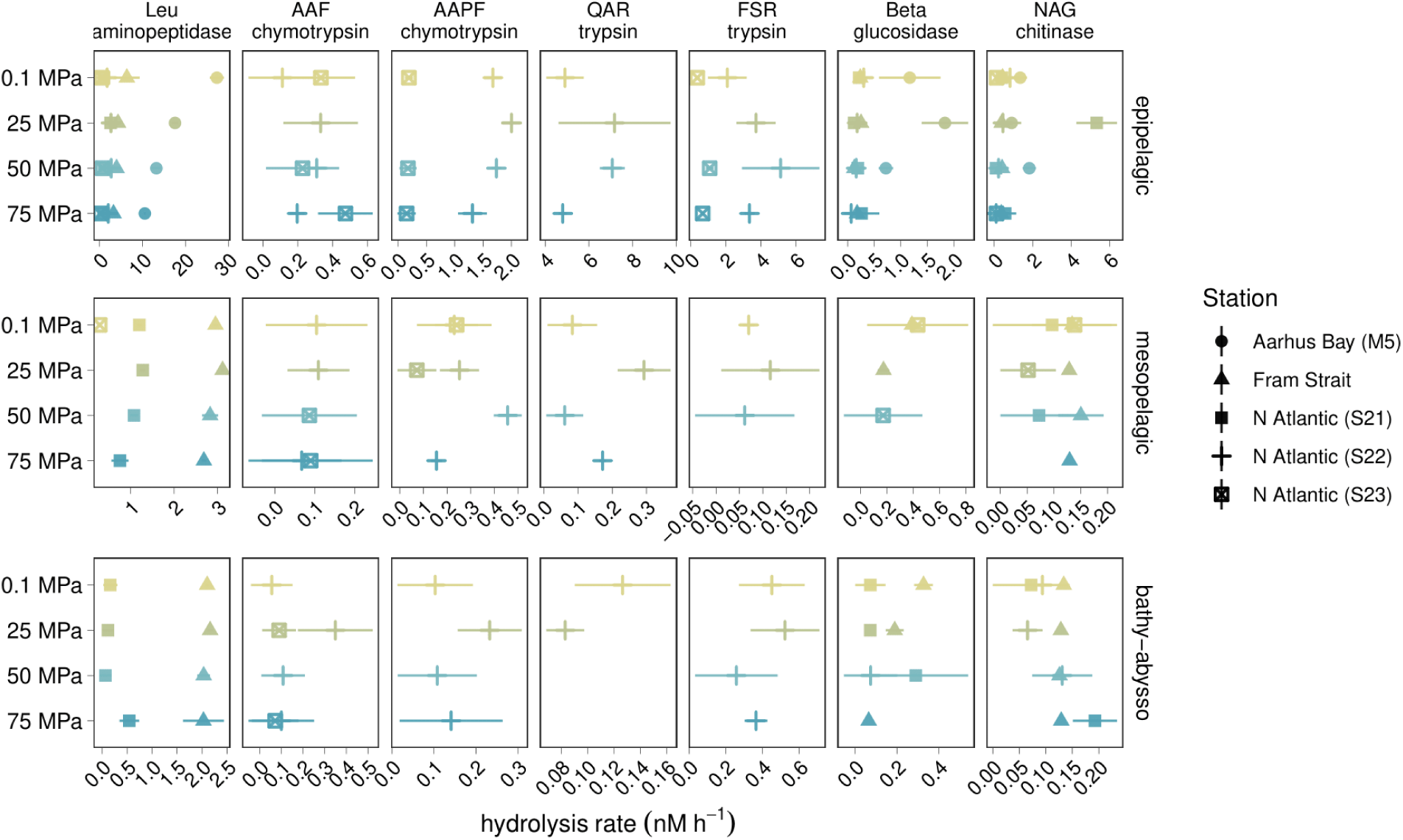
Enzyme hydrolysis rates for pelagic bacterial communities, separated by depth layer (e.g., epipelagic, mesopelagic, and bathypelagic to abyssopelagic); see Table 1 for specific depths. Error bars show the standard deviations of triplicates. Leu = leucine; AAF = alanine-alanine-phenylalanine; AAPF = alanine-alanine-proline-phenylalanine; QAR = Boc-glutamine-alanine-arginine; FSR = Phenylalanine-serine-arginine; NAG = N-acetyl-glucosaminide (substrate for chitinase).

### Pressure effects on cell-free versus cell-present enzyme activities

The extent to which pressure acts on extracellular enzymes or on the producer cells—and their capacity to produce more enzymes—has different ecological and biogeochemical implications. Whereas the former would indicate pressure effects on enzyme structure that affects hydrolysis rates—such as through increased hydration (Ohmae et al., 2013) or protein subunit dissociation (Ohmae et al., 2007) that changes the conformation and catalytic efficiency of enzymes—the latter suggests altered physiology and gene transcription that shifts enzyme production in response to pressure (Vezzi et al., 2005; Campanaro et al., 2012). Our measurements above reflect both of these factors. To identify the extent to which hydrostatic pressure can lead to physical-chemical effects on cell-free enzymes from natural assemblages at a select set of stations and depths (Table 1a), we compared cell-free (<0.2 µm) versus cell-present (≥0.2 µm) enzyme activities at different pressures. We incubated microbial communities with particulate organic matter (POM), and then separated the cell-associated from the freely released enzymes by filtering a portion of the incubation through a 0.2 µm filter. The rationale is that POM addition can greatly increase the enzyme activities of natural communities (Balmonte et al. 2019; Brown et al. 2022), likely yielding sufficient cell-free enzymes to carry out these measurements. Our marine POM sources were freshly collected live sargassum, and high molecular weight *Thalassiosira weissflogii* (diatom) biomass processed as previously described to remove low molecular weight organic matter (Balmonte et al., 2019); we additionally used one terrestrial POM source containing a mixture of dried plants (i.e., cereal grasses).

Overall, these comparisons show variable effects of pressure on cell-free vs. cell-present enzyme activities (Fig. 4a), ranging from substantial pressure reduction (e.g., NAG chitinase for North Atlantic, Stn 22, stimulated with sargassum and *T. weissflogii*), to relatively limited pressure inhibition (e.g. leucine aminopeptidase stimulated with *T. weissflogii* in North Atlantic, Stn. 22 and 23). Incubations in which only cell-free enzyme activities were measured also demonstrate similar variability (Supporting Information, Fig. S3-S4). Pressure reduction is observed for cell-free enzyme leucine aminopeptidase in the sargassum and *T. weissflogii* addition (Supporting Information, Fig. S3), and all enzymes tested with the cereal grass stimulation (Supporting Information, Fig. S4). Consequently, the relative contributions of cell-free vs. cell-present enzyme activities to total activities are also variable and show one of three trends with increasing pressure (Fig. 4b): (1) decreasing proportion of cell-free enzyme activities (e.g., beta-glucosidase stimulated with sargassum stimulation in North Atlantic Stn 22), (2) increasing proportion of cell-free enzyme activities (e.g., AAPF chymotrypsin throughout all of the POM stimulation experiments), or (3) relatively invariable proportion of cell-free enzyme activities (e.g., AAF chymotrypsin with *T. weissflogii* stimulation in North Atlantic Stn 23). Thus, we observe evidence that increasing pressure may reduce cell-free enzymes activities to a greater as well as to a lesser extent than cell-present enzyme activities, highlighting the complex pressure responses of a mixture of diverse cells and enzymes, consistent with observations among pelagic (Fig. 3) and benthic (Fig. 2) microbial communities.

**Figure 4.**
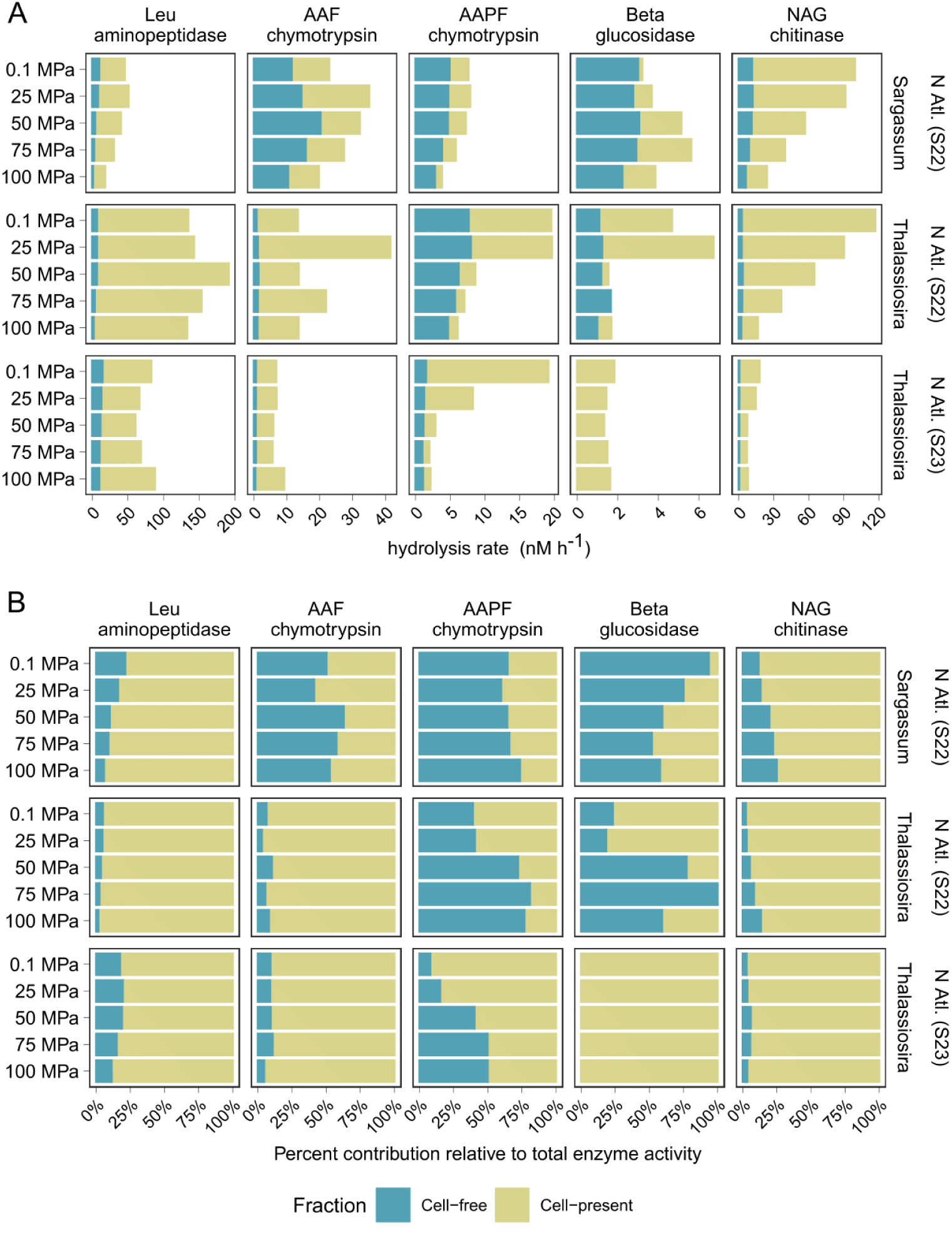
Hydrolysis rates and relative contributions of cell-free (<0.2 µm) vs. cell-present (bulk rates minus cell-free rates) pelagic enzyme activities following stimulation with sargassum or high molecular weight *Thalassiosira weissflogii* biomass at different stations and depths, shown either as stacked bar graphs (A) or filled bar graphs (B) to demonstrate the percent contribution of each fraction relative to total enzyme activities. Leu = leucine; AAF = alanine-alanine-phenylalanine; AAPF = alanine-alanine-proline-phenylalanine; NAG = N-acetyl-glucosaminide (substrate for chitinase). ATM = atmospheric pressure (0.1 MPa).

The pressure reduction of both cell-free (Supporting Information, Fig. S3-S4) and cell-present enzyme activities (Fig. 4) is consistent with prior studies (reviewed in Tamburini et al., 2013 and Picard and Daniel, 2023; Lloyd et al., *in revision*). Variability in pressure reduction is observed across depths, stations, and POM sources. For example, cereal grass stimulated the production of enzymes that are consistently and strongly reduced by pressure, whereas stimulation with sargassum and *T. weissflogii* did not. Presumably, the use of different POM sources stimulated distinct microbial taxa and or a distinct set of enzymes; thus, different pressure effects on a variety of cell-free enzymes may reflect their varying pressure sensitivities. These results are in accordance with findings that different types and concentrations of organic substrates can lead to differing pressure responses among microorganisms (Jannasch and Wirsen, 1982). In cases of substantially lowered catalytic activity of cell-free enzymes, leading to decreased relative contributions of cell-free enzymes at higher pressures, microorganisms would need to increase enzyme production to maintain the same level of function. Pressure-induced increased transcription of genes for enzymes (Vezzi et al., 2005) suggests that this strategy is feasible, but likely energetically expensive.

However, the observation that cell-free enzymes can maintain relatively robust functionality at high pressures is notable (i.e., >70% activity at high pressure relative to atmospheric pressure) (Fig. 4a; Supporting Information, Fig. S3). Such findings are most evident in AAF chymotrypsin and a few instances with AAPF chymotrypsin, β-glucosidase, and NAG chitinase (Supporting Information, Fig. S3). Prior studies of enzymes, such as the protease trypsin, from atmospheric pressure-adapted microorganisms have shown some similar pressure responses, with increased activation up to 40 MPa and a subsequent plateauing of catalytic efficiency at higher pressures (Groß et al., 1993). Hence, our findings suggest that even when produced by surface-originating microbial communities under atmospheric pressure conditions (e.g., stimulation of epipelagic microbial communities with sargassum), certain cell-free enzymes that hydrolyze the substrates tested here may continue to retain functionality up to hadal pressures. Whether these same organisms can still produce enzymes at high pressure is an open question. Strong pressure reduction of cell-present enzyme activities suggests increasingly low production of new enzymes at high pressure (Fig. 4b), consistent with prior studies that show severely stunted growth and respiration of bacterial cells at pressures >40 MPa (Bartlett and Marietou, 2014; Stief et al., 2021; Stief et al., 2023). However, the potential for some degree of pieozotolerance, even among surface-originating microbes, suggests the possibility of sustained enzyme production even with reduced efficiency (see *Biogeochemical and Ecological Implications*).

Findings of relatively high enzyme function at high pressures contrast with effects measured for polysaccharide hydrolase activities in experiments with coastal seawater, using the same high molecular weight *Thalassiosira* biomass. In that study, increased hydrostatic pressure resulted in complete inhibition of cell-free enzymatic activities (Lloyd et al., *in revision*). However, the polysaccharide hydrolyzing enzymes whose activities were measured in that study are very different from the peptidase enzymes whose activities were measured in the current study. In particular, polysaccharide hydrolases are very substrate-structure selective compared to peptidases (Lapebie et al., 2019), and there is a far lower possibility that a polysaccharide hydrolase will also act on a non-target substrate. Thus, polysaccharide hydrolysis is a very specific enzymatic process, and the distribution of specific polysaccharide hydrolase enzymes among microorganisms is likely far narrower than the distribution of peptidases and exo-acting glucosidases measured here. For polysaccharide hydrolases, a community might have relatively little functional redundancy in specific activities: pressure inhibition of a narrow range of enzymes would not be compensated by the availability of related enzymes that may also exhibit activity against a given substrate. Differences in enzymatic structure and substrate specificity, as well as the range and pressure sensitivities of the microorganisms that produce the hydrolytic enzymes likely account for the varying pressure effects observed here and in other studies (reviewed in Tamburini et al., 2013 and Picard and Daniel, 2023; Lloyd et al., *in revision*).

### Biogeochemical and ecological implications

The pressure responses of complex pelagic and benthic microbial communities are variable (Fig. 2-4), but the most common response is pressure reduction, even for a moderate piezophile exposed to pressures greater than its optimum (Fig. 1). These findings indicate that microorganisms would have reduced enzyme catalytic efficiency and overall reduced contribution to carbon degradation at higher pressures. In certain circumstances, severe inhibition of enzyme activities may well constitute a roadblock for initiating carbon remineralization. Polysaccharide degradation, for example, may be fully inhibited, as shown in a recent study (Lloyd et al., *in revision*). In such a situation, a lack of hydrolysis would limit the flow of carbohydrate substrates into a cell, restricting the carbon and energy available for other pressure-sensitive cellular processes, including microbial biomass production (Amano et al., 2022) and aerobic respiration (Stief et al., 2021; Stief et al., 2023; Franco-Cisterna et al., 2024).

Instances of less pronounced pressure reduction or even pressure resistance, however, are notable exceptions to the widely observed pressure reduction. Less severe pressure reduction of enzyme activities by benthic microbial communities (Fig. 2) leads to high measurable rates even at 75 MPa (Supporting Information, Fig. S2). The greater diversity and wider range of enzymatic capabilities of benthic versus pelagic microbial communities (Teske et al. 2011) increases the likelihood that some enzymes—or some organisms that can produce the enzymes— can withstand high hydrostatic pressures. In addition, cases of much lower pressure sensitivity are evident among cell-free and cell-present enzyme activities (Fig. 4, Supporting Information, Fig. S3-S4). Thus, cell-free enzymes and perhaps some microbial cells, such as those that sink associated with marine snow, may continue to transform particulate matter and high molecular weight dissolved organic matter to low molecular weight fractions even at high pressures.

The range and limits of enzyme activities at high pressures also have other ecological implications. Maintaining the ability to enzymatically hydrolyze organic matter up to hadal pressures may indicate that a subset of surface ocean-originating microbial taxa, at the very least, may be piezotolerant. Transcripts belonging to surface-originating microorganisms recovered in the deep bathypelagic (Poff et al., 2021) additionally provide evidence that surface microbes can maintain functionality in the deep, at least for a finite period. Piezotolerance could be particularly advantageous to organisms that sink associated with particles, especially under conditions of rapid carbon export that occur through particle injection pumps (Boyd et al., 2019; Poff et al., 2021), and as long as a subset of their populations return to shallow waters and as long as this trait does not compromise their competitiveness in shallow waters. Recent estimates that ca. 85% of bathypelagic microbial communities are piezotolerant, but only ca. 5% are piezophiles, may in part be related to the constant supply of piezotolerant organisms from surface waters that can seed deep water communities (Amano et al., 2022; Glud and Schauberger, 2024). Potentially widespread piezotolerance suggests that other microbially-mediated biogeochemical processes— including the production of diverse hydrolytic enzymes whose activities were not measured in the current study—may also proceed to an extent despite increased hydrostatic pressures. Future efforts to characterize the pressure regulation of other enzymatic activities, their molecular mechanisms, and the microbial taxa involved will provide a more comprehensive understanding of biogeochemical transitions that occur from the surface to the interior of the oceans.

## Acknowledgements

We are grateful to the many people who assisted with sample collection and experimental setup for this study. We thank the members of the HADAL Center and those at University of Southern Denmark, especially Anni Glud, Feiyang Gu, Morten Kieler, Mikkel Nielsen, Clemens Schauberger, Peter Stief, Sachia Traving, Bo Thamdrup, and Frank Wenzhofer, for field, technical, and experimental assistance. We thank the captain and crew of R/V *Aurora* during the 2021 Aarhus Bay cruise; the captain and crew of R/V *Polarstern* cruise during the 2021 Fram Strait (HAUSGARTEN) cruise PS126; Don Canfield and the captain and crew of his 2021 Skagerrak Strait cruise; and the captain, crew, and scientific party of R/V *Endeavor* EN683 North Atlantic cruise. This project was funded by the U.S. National Science Foundation (OCE-2022952 and OCE-2241720 to CA, and OCE-2241721 to JPB) and the Danish Center for Hadal Research (HADAL, Grant No. DNRF145 to RNG).

## Data Availability Statement

Data and metadata will be made available in the Biological and Chemical Oceanography Data Management Office (BCO-DMO) Data Repository.

