## Supplementary Information for "Effects of increasing hydrostatic pressures on marine microbial enzymatic activities"

**Running head**: Pressure effects on enzyme activities

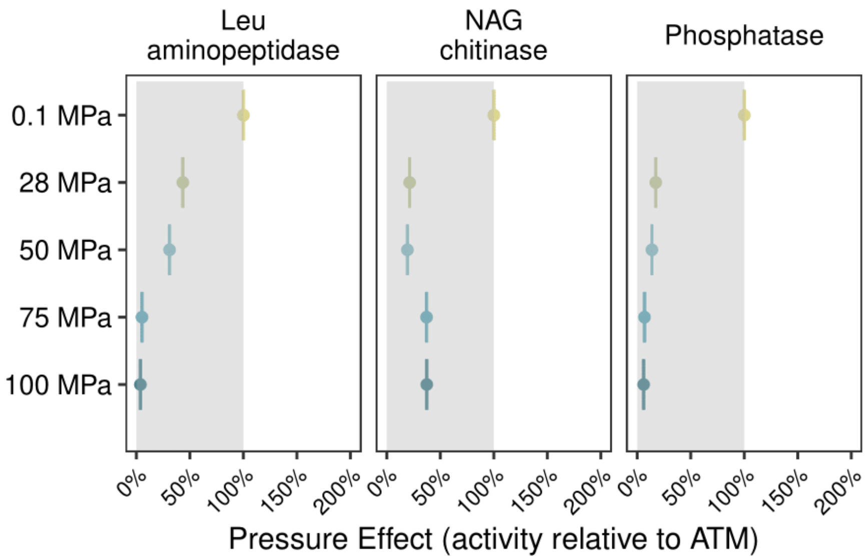


**Figure S1.** Pressure effects on the cell-specific enzyme activities of *Photobacterium profundum* SS9. The shaded box indicates the region of pressure reduction. Leu = leucine; NAG = N-acetyl-glucosaminide (substrate for chitinase). ATM = atmospheric pressure (0.1 MPa)

**Supplementary text.** To analyze *P. profundum* SS9 cell abundances, samples were fixed with glutaraldehyde to a final concentration of 1% and stored at -20 °C until processing using a BD FACSCanto™ II flow cytometer. SYBR Green I was used to stain the cells prior to analysis on the flow cytometer. BD Trucount™ beads were used to normalize the detected cell counts, and the flow cytometry data was processed using the program Flowing Software 2.

Cell counts after 24 hrs of incubation at each pressure level are as follows: 11.4 x$10^6$ cells $mL^{-1}$ (0.1 MPa), 106.1 x$10^6$ cells $mL^{-1}$ (28 MPa), 116.9 x$10^6$ cells $mL^{-1}$ (50 MPa), 18.1 x$10^6$ cells $mL^{-1}$ (75 MPa), and 16.2 x$10^6$ cells $mL^{-1}$ (100 MPa).

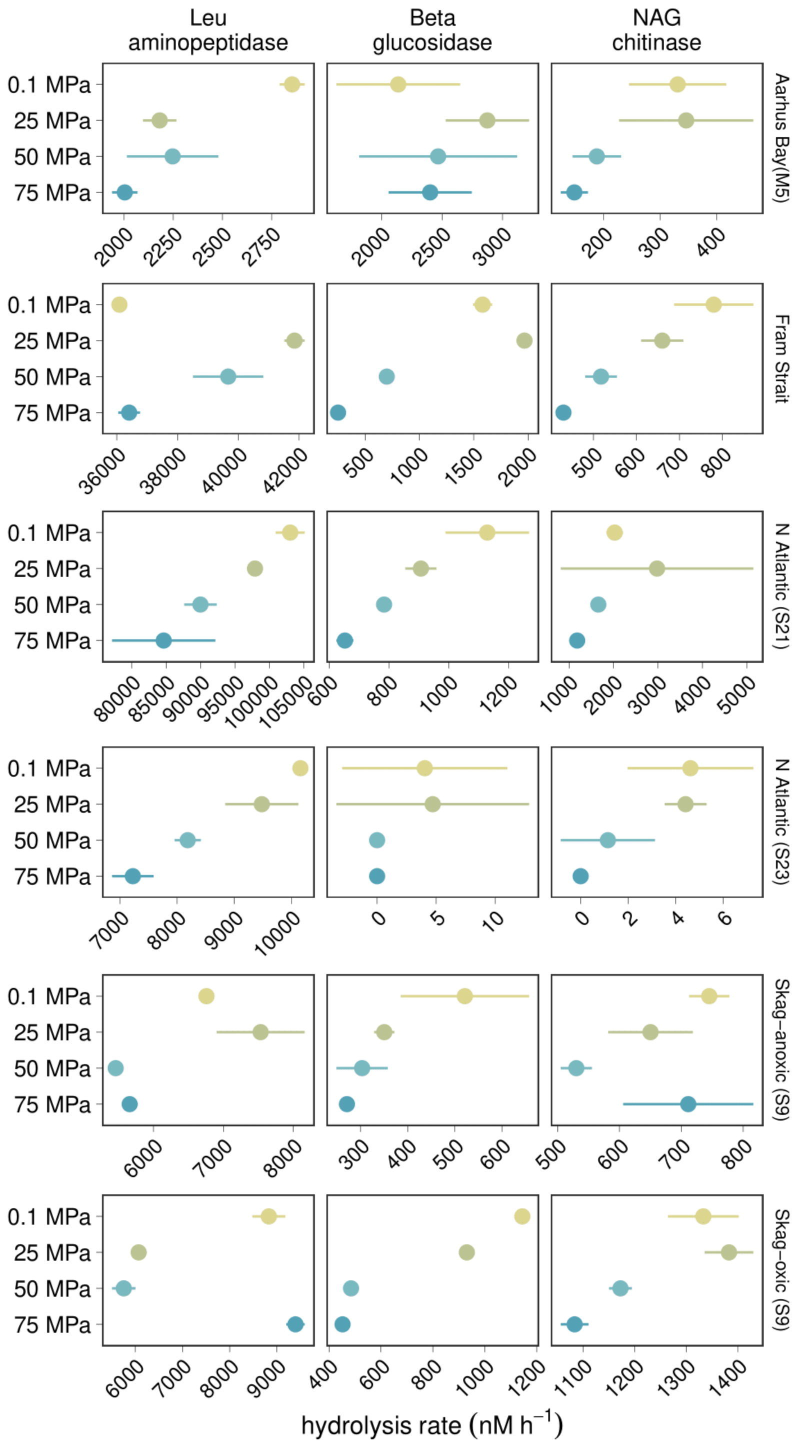


**Figure S2.** Enzymatic hydrolysis rates for sediment bacterial communities. Error bars show the standard deviations of triplicates. Leu = leucine; NAG = N-acetyl-glucosaminide (substrate for chitinase). Skag-anoxic = Skagerrak Strait (S9) anoxic; Skag-oxic = Skagerrak Strait (S9) oxic.

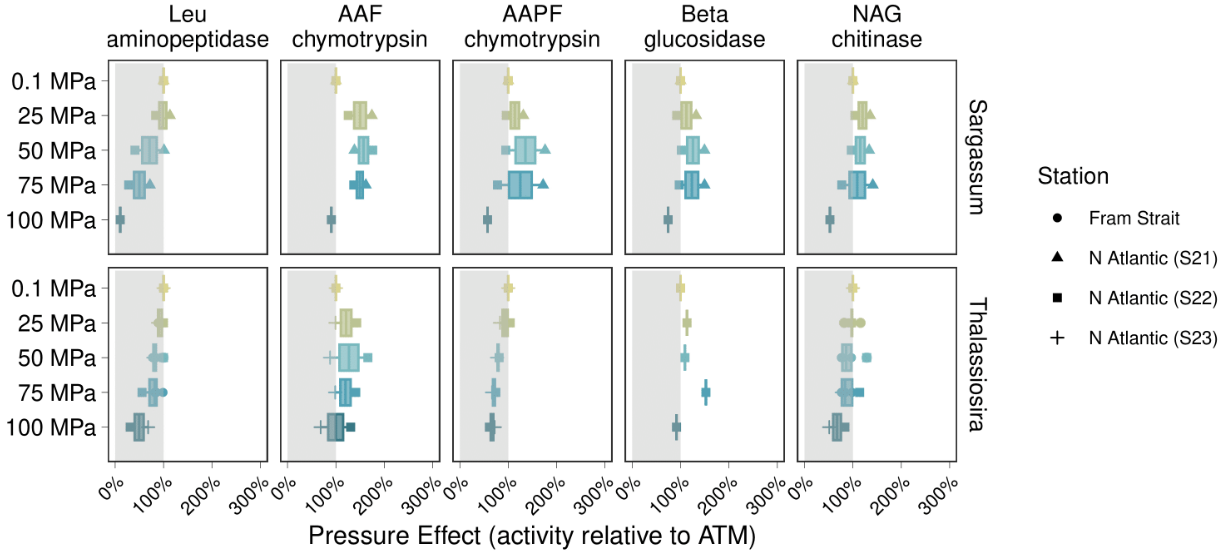


**Figure S3.** Pressure effects on cell-free (<0.2 µm) enzymes produced by pelagic microbial communities from the Fram Strait and North Atlantic Ocean following stimulation with sargassum (top) or high molecular weight *Thalassiosira weissflogii* biomass (bottom). The shaded box indicates the region of pressure reduction. Leu = leucine; AAF = alanine-alanine-phenylalanine; AAPF = alanine-alanine-proline-phenylalanine; NAG = N-acetyl-glucosaminide (substrate for chitinase). ATM = atmospheric pressure (0.1 MPa).

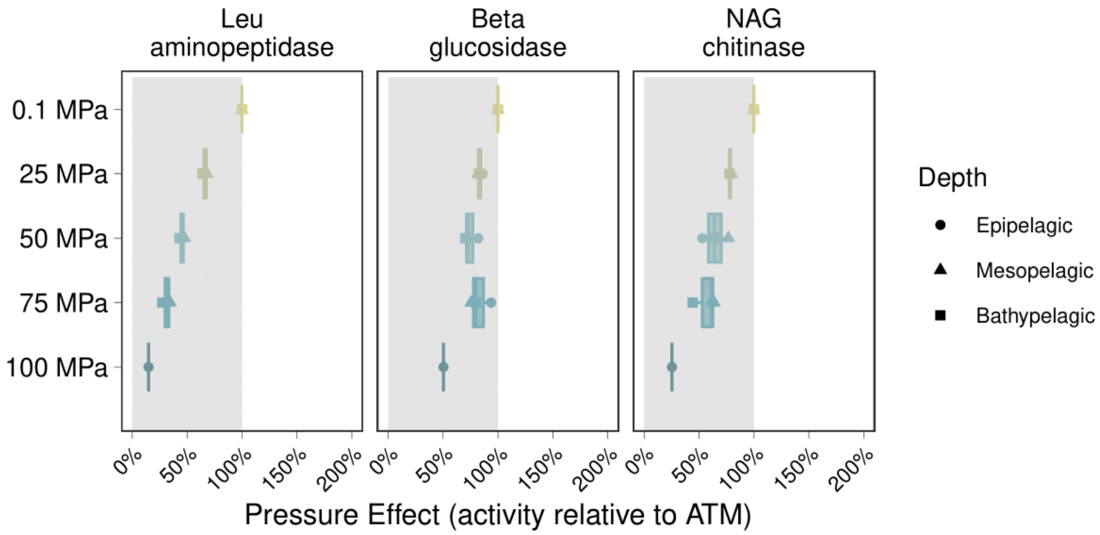


**Figure S4.** Pressure effects on cell-free enzymes (<0.2 μm) produced by pelagic microbial communities from [[*location*]] following stimulation with dried cereal grass leaves, a representative terrestrial source of particulate organic matter. The shaded box indicates the region of pressure reduction. Leu = leucine; NAG = N-acetyl-glucosaminide (substrate for chitinase). ATM = atmospheric pressure (0.1 MPa)

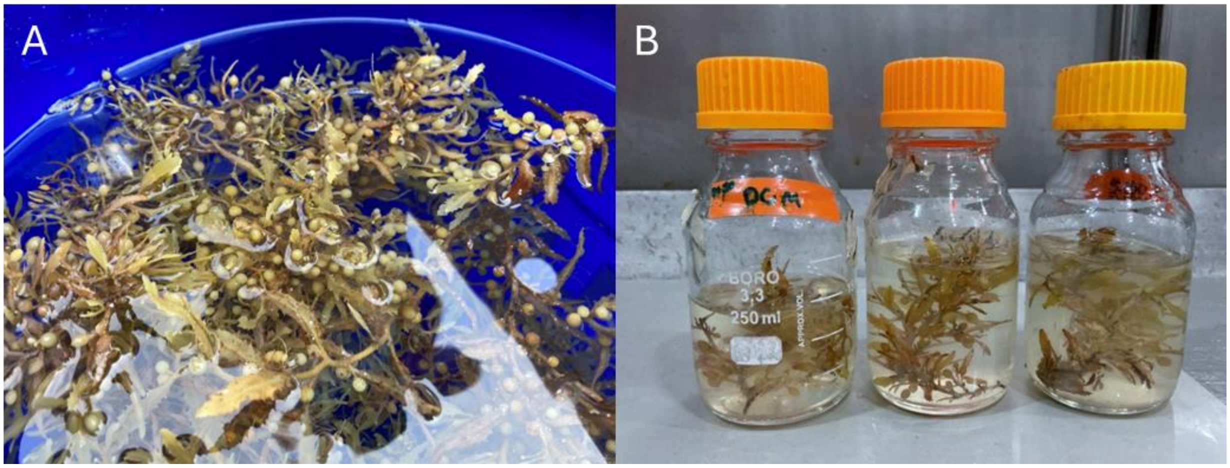


**Figure S5.** Freshly collected sargassum prior to incubation with seawater (A), and sargassum after incubation with seawater from different depths to stimulate enzyme production (B). Photo to the right (B) was taken after setup of pressure experiments. Incubations to stimulate enzyme production included the sargassum pieces (B) and 250 mL of water per depth.
